# Prediction of context-specific regulatory programs and pathways using interpretable deep learning

**DOI:** 10.1101/2024.11.06.622202

**Authors:** Daria Doncevic, Carlos Ramirez Alvarez, Albert Li, Youcheng Zhang, Anna von Bachmann, Kasimir Noack, Carl Herrmann

## Abstract

Variational autoencoders (VAEs) are being widely adopted for the analysis of single-cell RNA sequencing (scRNA-seq) data. As with any non-linear models, however, they lack interpretability, which is a crucial aspect in the biomedical field where researchers want to be able to trust their model predictions. Our previously developed OntoVAE model addressed this issue by integrating biological ontologies in the decoder, which made the neuronal activations correspond to pathway activities. However, when multiple covariates are present, disentangling their relative contributions is challenging. To address this limitation, we developed COBRA, a VAE tool that combines the interpretable decoder part of OntoVAE with an adversarial approach that separates covariate effects in the latent space. In this work, we demonstrate the use of COBRA on two different scRNA-seq datasets in different contexts. We applied the tool to an interferon stimulated mouse dataset to separate the effects of celltype and treatment on transcription factors and biological pathways. We furthermore showed how COBRA can be used to predict the state of unseen celltypes.

## 1 Introduction

Deep learning models such as Variational Autoencoders (VAEs) are now widely used to analyze single-cell data for different purposes such as data denoising, clustering, data integration, and perturbation analyses [1–3]. VAEs perform dimensionality reduction of the data and capture the essence of the data in their latent space [4]. The latent space embedding is then used for subsequent tasks such as clustering and cell type annotation [5]. However, as with any non-linear model, the latent space of VAEs is not directly interpretable, as the contributions of the different features cannot be easily calculated [6]. Thus, the field of interpretable deep learning has emerged that addresses this issue [7]. There are two main approaches taken by the field: post-hoc and intrinsically interpretable methods. Post-hoc methods are applied on an already trained model. LIME for example uses local approximations with linear models [8], while SHAP considers possible combinations of input features to extract single contributions [9]. Intrinsically interpretable models adapt the model structure to achieve interpretability by design. Some methods incorporate biological information directly into the model as prior. VEGA, for example, couples a sparse, one-layer decoder to a fully connected encoder. Each neuron in VEGAs latent space corresponds to a biological entity – a transcription factor (TF) or a pathway – and is only connected to annotated genes in the output layer, thus resulting in a sparse decoder. Thus, the latent space dimensions of VEGA are directly interpretable as TF or pathway activities [10]. Other models incorporate hierarchical biological information, such as DCell, a neural network that is structured according to Gene Ontology (GO) subsets [11], and our own OntoVAE model, a VAE that combines a standard encoder with a multi-layer sparse decoder that can accommodate any biological ontology [12]. Similar to VEGA, the activations of the neurons in the decoder of OntoVAE correspond to pathway activities. OntoVAE can also be used in the context of perturbation modeling, to predict for example the outcome of genetic perturbations or drug treatments. In this type of analysis of heterogeneous single-cell datasets, the cell type can be a strong confounder which masks the effect of interest. To address this, we expanded our OntoVAE model with an adversarial approach that was previously introduced in the CPA model [13] and encourages the disentanglement of different covariates in the latent space. The new model, COBRA (COvariate Biological Regulatory network Autoencoder), allows dissecting the contributions of different covariates while maintaining the interpretability in the decoder.

## 2 Results

### 2.1 Architecture of COBRA

COBRA combines the interpretable decoder part of OntoVAE with an adversarial approach that allows for a disentanglement of specified covariates in the latent space of the model (**Figure 1a**). The details of this adversarial approach have already been described in [13] and are also given in the *Methods* section. In brief, auxiliary classifiers for each covariate are attached to the latent space of the model and encourage the learning of a so-called basal state, where any covariate influence is removed. Simultaneously, embedding layers learn the embeddings of the different covariates, and these then get added to the basal state. During model training, the sum of the basal state and the covariate embeddings is passed to the interpretable decoder for reconstruction. The model is trained end-to-end, alternatively updating the weights of VAE and embedding layers, and of the auxiliary classifiers. As with OntoVAE, COBRA allows for incorporation of any biological network, such as gene regulatory networks (GRNs) or ontologies. Such networks can be composed of a single layer representing TFs or Reactome pathways but can also be multi-layered in the case of a hierarchical ontology. Each decoder node (TF, Reactome pathway or ontology term) is connected to the corresponding genes in the output layer. Importantly, once a model is trained, TF or pathway activities can be retrieved for each covariate or view separately. Note that the terms ‘covariate’ and ‘view’ are used interchangeably throughout this manuscript, and ‘view’ can denote any combination of the observed covariates.

**Fig. 1.**
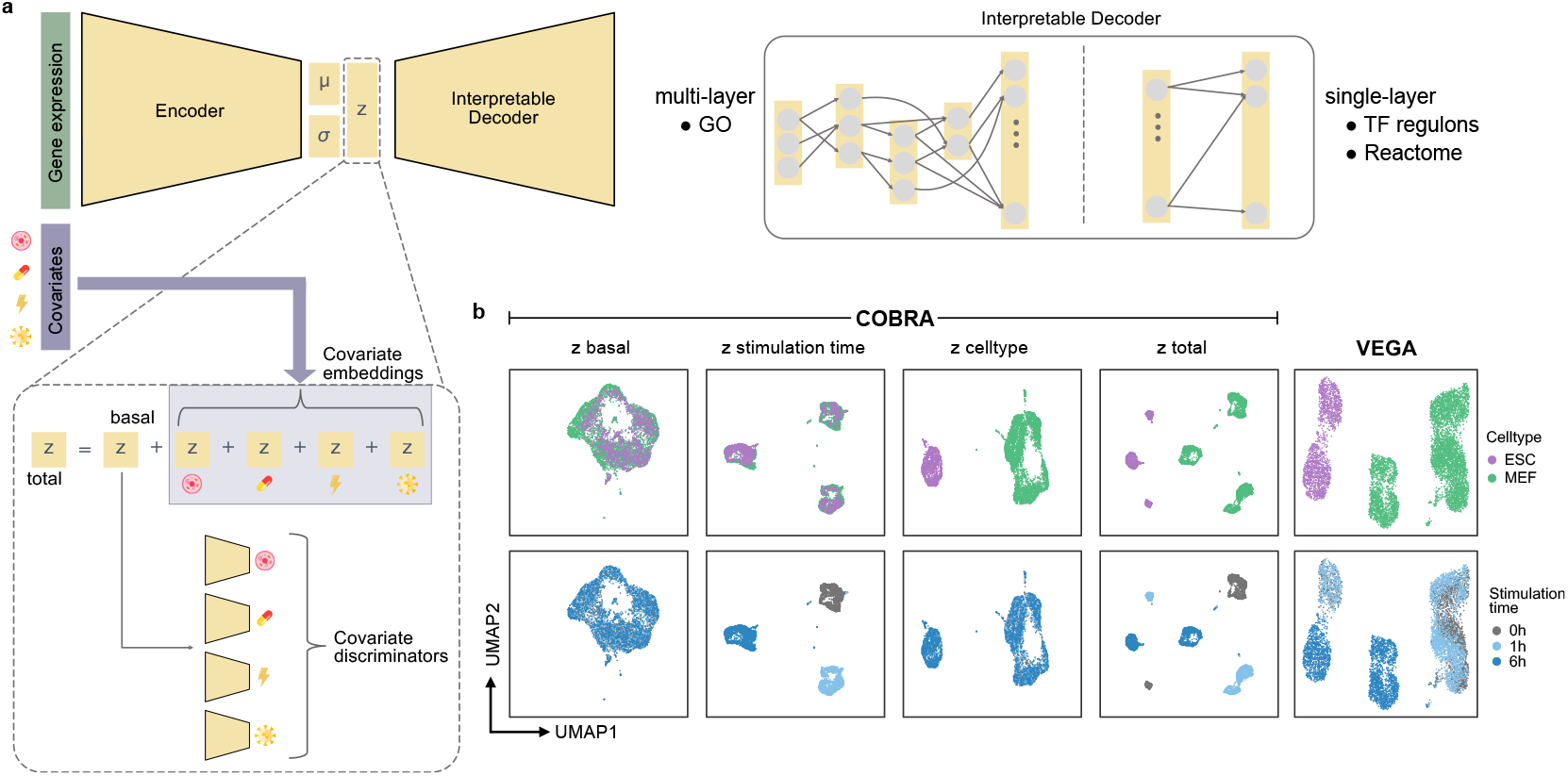
COBRA combines disentanglement of covariates with interpretability. (a) Schematic representation of COBRA. A standard encoder is coupled to a sparse decoder which reflects a biological regulatory network, that can be multi-layered or single-layered. Covariate effects are separated in the latent space of the model. Auxiliary classifiers remove covariate information and achieve a mixing of cells (basal state). The covariate information is learned separately through embedding layers and then added to the basal state. The sum (total) is then fed into the decoder. (b) COBRA was applied on a mouse interferon dataset using the collecTRI TF regulons as prior. Displayed are the UMAPs of the latent space embeddings for the different views of COBRA – basal, stimulation time, celltype, and total – and for VEGA, which was applied on the same dataset for comparison.

### 2.2 COBRA disentangles celltype and interferon treatment effect

We trained COBRA on a previously published scRNA-seq dataset of mouse embryonic stem cells (ESCs) and mouse embryonic fibroblasts (MEFs) that had been stimulated with interferon (IFN) beta for 0h, 1h, or 6 hours [14]. As a prior in the decoder, we used the mouse TF regulons from the collecTRI database [15]. For comparison, we trained VEGA [10] with the same prior. We then extracted the latent space embeddings from both trained models. In the case of VEGA, this corresponds to a single embedding with the dimension equal to the number of TFs. In the case of COBRA, we obtain four different embeddings: one for the basal state, one for the sum of basal state and celltype covariate, one for the sum of basal state and IFN stimulation covariate, and one for the total state (the sum of the basal state and both covariates). We calculated the UMAP on all embeddings (**Figure 1b**). For COBRA, we can observe that cells are well mixed in the basal view, but cluster according to celltype in the celltype view, cluster according to stimulation time in the stimulation time view, and form separate clusters per covariate combination in the total view. For VEGA, the cells are mainly clustering by celltype, but also by IFN stimulation time.

We then set out to investigate the TF activities for the two models. We selected important TFs by calculating the variance of their activities as reported in the IFN stimulation view of COBRA and then taking the top 60 TFs, which are represented in a heatmap in **Figure 2a**. We find many TFs that are central in interferon response, such as IRF7, IRF2, IRF1, STAT1, IRF3, IRF9, IRF4, IRF5, and STAT2 (all highlighted in the heatmap). VEGA also detects a change in the activities of these TFs upon stimulation. However, COBRA captures effects that are missed by VEGA. We illustrate this with NANOG, a TF that is crucial for the self-renewal and pluripotency of stem cells [16]. Accordingly, both the celltype view of COBRA and VEGA show a high activity of NANOG in the ESCs compared to MEFs. However, COBRA additionally reveals activation of NANOG upon IFN treatment (**Figure 2b**). This interferon dependent increase is confirmed in several studies, for example using a sarcoma cell line [17] or after transfection with dsRNA [18]. We hypothesize that the strong celltype specific effect overshadows the potential response to IFN of NANOG in this dataset.

**Fig. 2.**
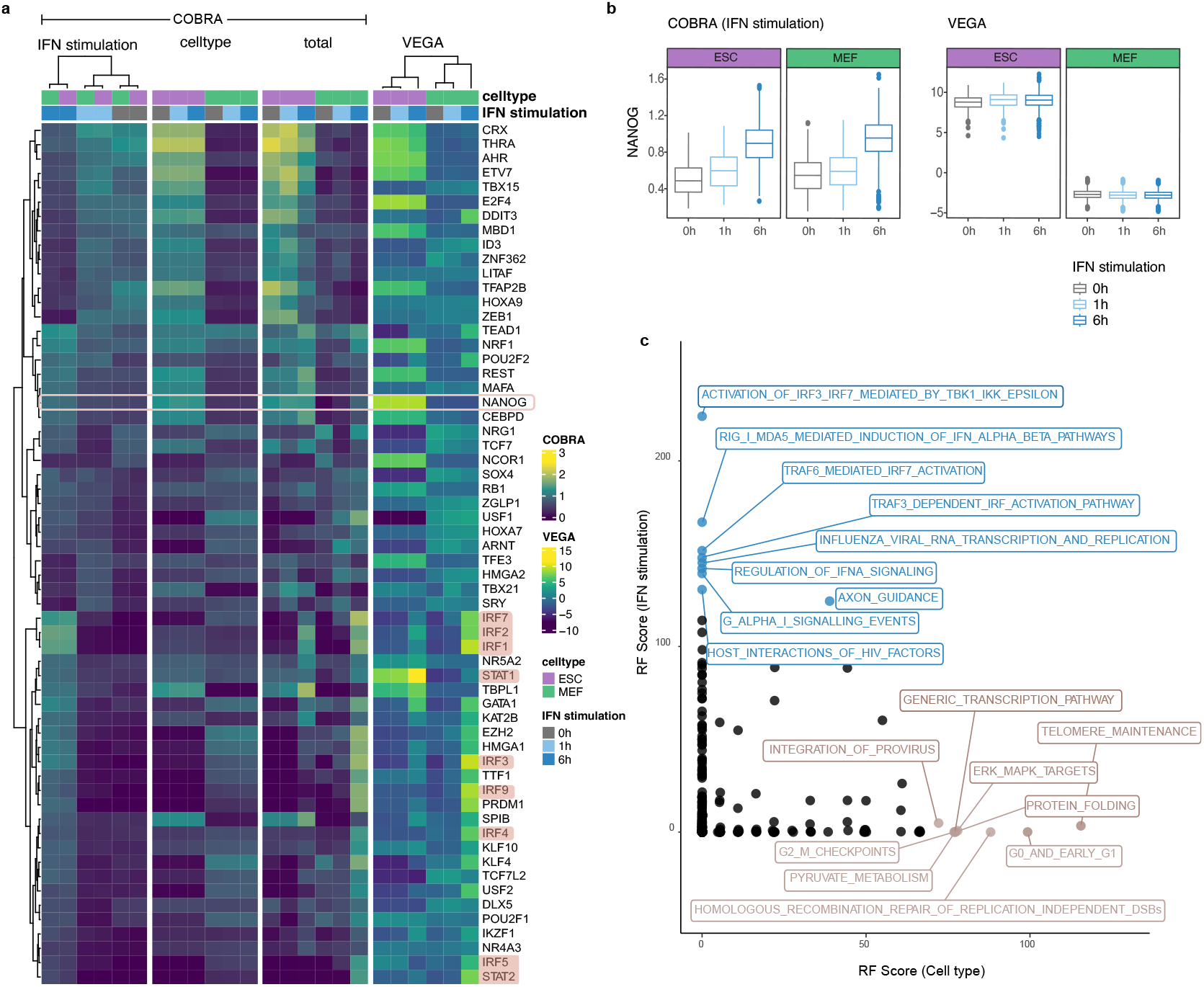
COBRA disentangles celltype from stimulation effect in mouse interferon dataset. (a) Heatmap comparing TF activities between the different views of COBRA and VEGA using gene regulatory networks as biological prior. The top 60 TFs with the highest variance aggregated by celltype and stimulation time are displayed. Highlighted are TFs that are known to be involved in IFN response (STAT1, STAT2, IRFs) as well as NANOG (b) Boxplots compare the TF activities of NANOG between the IFN stimulation view of COBRA and VEGA. (c) Random forest classification using the pathway activities retrieved from the IFN stimulation and the celltype view, respectively, as features. Feature importances were then extracted for all pathways for both classification analyses and plotted agains each other. Labelled are the top 9 pathways for IFN stimulation (blue) and celltype (brown), respectively.

We then trained COBRA again on the same dataset, now using Reactome pathways as a prior, and extracted the pathway activities for the different views. We performed a random forest classification to classify IFN stimulation timepoints based on the pathway activities from the stimulation time view, and celltype based on the pathway activities from the celltype view, respectively. The feature importances for both classification analyses are plotted against each other in **Figure 2c**. Most pathways were found to have a high importance in classifying one of the two covariates, and low importance for the other covariate, respectively. For the classification of celltype, we found many pathways that are related to stemness, such as ‘Telomere maintenance’, ‘G0 and early G1’, ‘G2M checkpoints’, and ‘Homologous recombination repair of replication independent double strand breaks’. Among pathways that are associated with IFN stimulation time, we found pathways related with IFN or IRFs, such as ‘Activation of IRF3 IRF7 mediated by TBK1 IKK epsilon’, ‘RIG1 MDA5 mediated induction of IFN alpha beta pathways’, and ‘TRAF6 mediated IRF7 activation’. Taken together, these results indicate that COBRA effectively decouples celltype from treatment effect in the mouse dataset, and captures the relevant pathways for both covariates.

### 2.3 COBRA isolates timepoint effect in the developing adrenal medulla

Next, we applied COBRA on an scRNA-seq dataset of the developing human adrenal medulla, sampled at different timepoints post-conception [19]. The lineage trajectories are visualized in a UMAP that was computed on the expression data (**Figure 3a**): Schwann cell precursors (SCPs) differentiate over intermediate celltypes into two separate lineages: neuroblasts and chromaffin cells. Celltype abundances are correlated with the developmental timepoint measured in post-conception weeks (pcw), with a larger amount of Late SCPs, Late neuroblasts, and Late chromaffin cells in the later timepoints 11pcw, 14pcw, and 17pcw (**Figure 3b**).

**Fig. 3.**
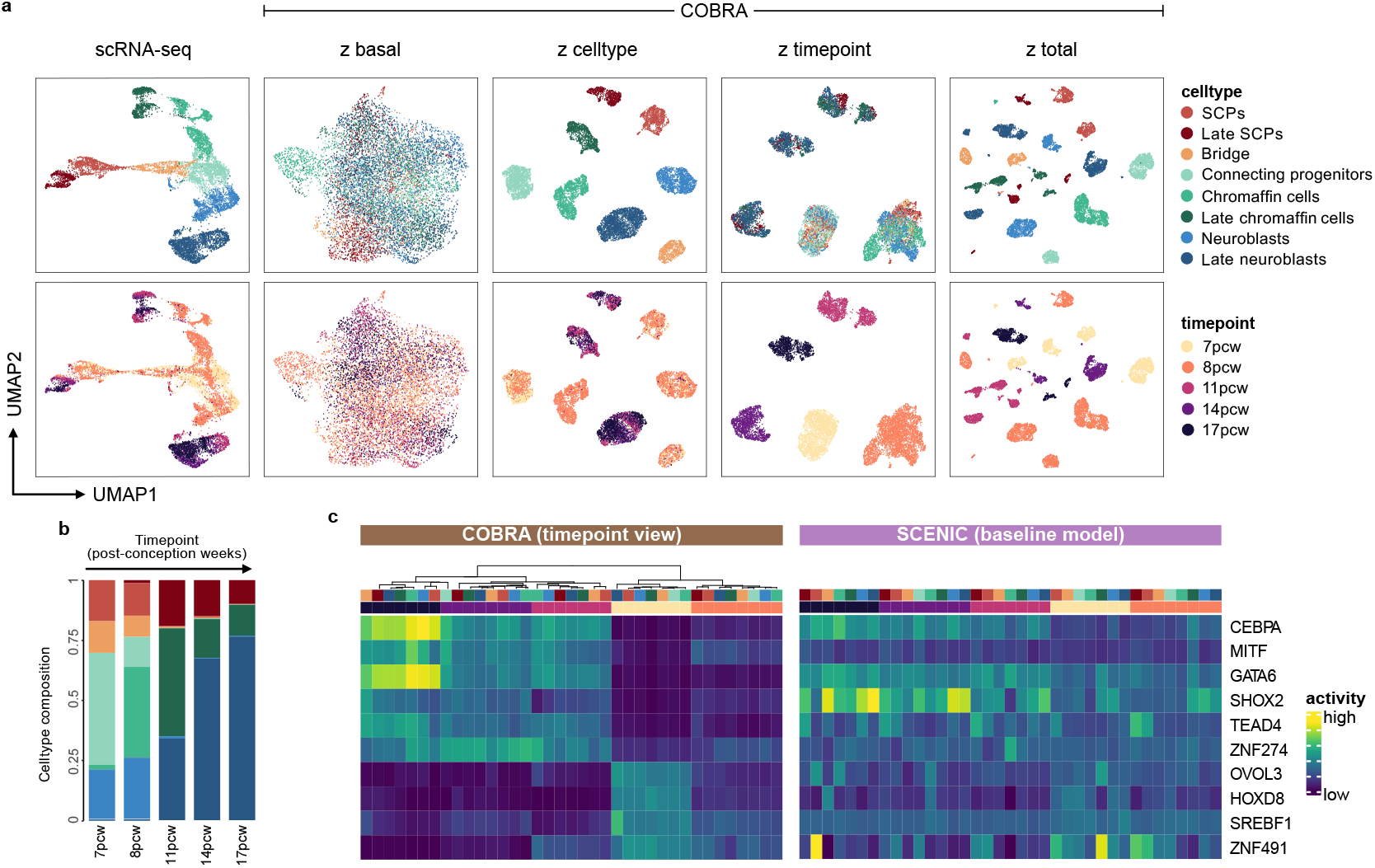
COBRA can disentangle the timepoint from the celltype effect. COBRA was trained on all cells from the adrenal medulla dataset [19] using TF regulons computed with SCENIC as prior. (a) UMAPs on the scRNA-seq data and the different latent space views of COBRA (z basal, z celltype, z timepoint and, z total). Cells are colored by celltype (top) or timepoint (bottom). (b) Celltype composition in the different timepoints. (c) The 10 TFs with the strongest correlation between their activity in the timepoint view and the timepoint were selected. Heatmaps display the activity of those 10 TFs in the timepoint view of COBRA and as calculated by SCENIC (baseline model).

In their work, Jansky et al. identified TFs that are important for the different celltypes [19]. We wanted to see whether COBRA could identify TFs that are related with cellular differentiation independently of the celltype. For this purpose, we trained the model with the covariates celltype and timepoint on all cells (full model), using the TF regulons computed by SCENIC as a prior. The UMAPs that were calculated on the latent space embeddings of the different views are displayed in **Figure 3a**. As expected, cells are mixed in the basal state, and cluster according to the respective covariate in the other views.

We then extracted the TF activities from the timepoint view, and computed their Pearson correlations with timepoint. The top 10 TFs that were selected this way are displayed in a heatmap (**Figure 3c**, left panel). CEBPA, MITF, GATA6, SHOX2, TEAD4, and ZNF274 have a positive correlation with timepoint for COBRA, and OVOL3, HOXD8, SREBF1, and ZNF491 a negative correlation with timepoint. CEBPA is known to regulate myeloid cell differentiation [20], MITF drives melanocyte differentiation [21], GATA6 leads to differentiation [22], SHOX2 is essential during heart development [23], TEAD4 plays an important role in trophectoderm differentiation [24], OVOL3 is involved in epidermal cell differentiation, and SREBF1 promotes adipocyte differentiation [25]. Furthermore, it is known that HOX genes influence stem cell differentiation during development [26]. Hence, the identified TFs are strongly enriched for developmental regulators. For comparison, we also show the activity of these TFs as calculated by SCENIC (**Figure 3c**, right panel). In the SCENIC analysis, the TFs MITF, TEAD4, ZNF274 and OVOL3 are not significantly associated with timepoint (p-value > 0.5), and the remaining TFs do not reveal as clear a timepoint associated pattern as with COBRA. This highlights the benefit of decoupling the celltype effect from timepoint to reveal stronger associations.

### 2.4 COBRA can predict the state of unseen celltypes

We were then interested to evaluate whether COBRA could make predictions about a celltype of the adrenal medulla dataset not encountered during training (out-of-distribution (ood) situation). For this purpose, we trained the model two more times: once on all cells except Late neuroblasts (nb-model), and once on all cells except Late chromaffin cells (chrom-model). For this analysis, we exploited the fact that Late neuroblasts and Late chromaffin cells can be considered as Neuroblasts and Chromaffin cells from later timepoints, respectively, and thus can use their covariate embedding when being passed through a trained model.

First, we wanted to check how well the nb-model and the chrom-model could reproduce the TF activities from the timepoint view of COBRA, given the absence of a substantial part of cells from later timepoints. We computed the correlations between TF activities and timepoint for the two ood models, and compared them against the correlations of the full model (**Figure 4a** for nb-model, **Figure 4b** for chrom-model). We observe a strong agreement for both models (nb-model: R=0.86, chrom-model: R=0.78). Some TFs were predicted to have an opposite correlation with timepoint compared to the full-model (dots colored in red). However, these TFs have lower correlations with timepoint in general (they are clustered more close towards the origin in the scatterplot), and are thus not as important to describe this covariate.

**Fig. 4.**
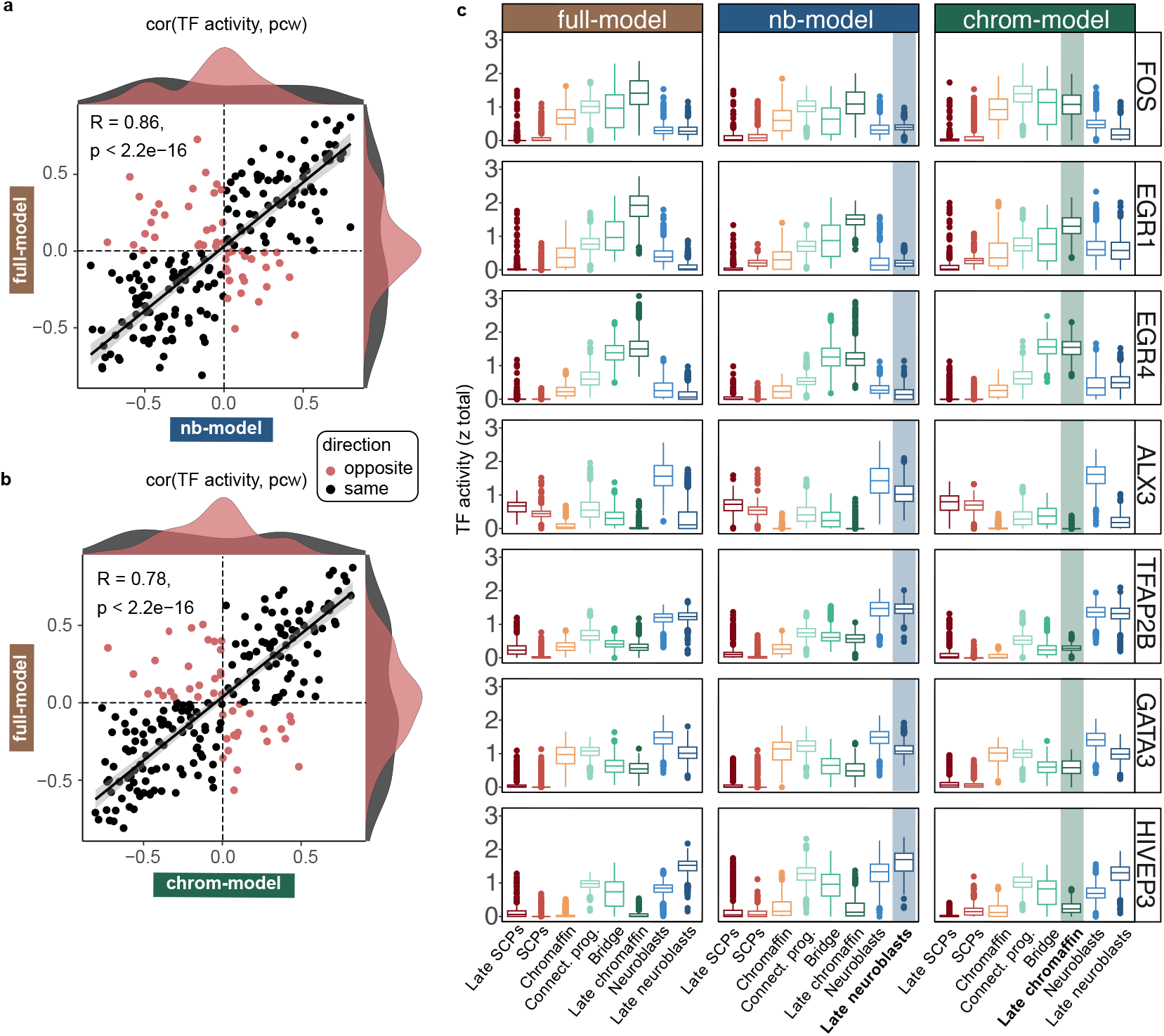
COBRA can predict late celltype effects. COBRA was trained on all cells from the adrenal medulla dataset excluding the Late neuroblasts (nb-model) or excluding the Late chromaffin cells (chrom-model). (a,b) Scatterplots show how well the TF activities in the timepoint views of the nb-model (a) and the chrom-model (b) agree with the full model. Each dot represents a TF, displayed are the Pearson correlations of TF activities with timepoint. TFs for which ood model and full model have the same +/sign are colored in black, TFs with opposite signs are colored in red. (c) Box plots display how well the nb-model and the chrom-model can predict the TF activities for the unseen celltype (color highlighted) for a selection of seven TFs important for either Neuroblasts or Chromaffin cells. TF activities were taken from the total view of COBRA.

Finally, we set out to investigate if COBRA could predict the TF activities of the celltypes that were not encountered during training. For this analysis, we selected TFs that were identified by Jansky et al. to be important for neuroblasts (ALX3, TFAP2B, GATA3, HIVEP3), or chromaffin cells (FOS, EGR1, EGR4). We then compared the activities for these TFs from the total view of COBRA between the two ood models and the full model. The results are displayed in **Figure 4c**, the predicted celltype is highlighted in color. We can see that the chrom-model captures the same trend between Chromaffin cells and Late chromaffin cells for EGR1 and EGR4 as the full model and the nb-model, which have both encountered Late chromaffin cells during training. For FOS, the chrom-model predicts no change, while its activity increases in the full-model. The nb-model captures the same trend between neuroblasts and Late neuroblasts as the full model and the chrom-model for all four neuroblast related TFs, regardless of the direction of the trend. Taken together, these results confirm that model training is robust, and that COBRA can make predictions about celltypes that were not encountered during training.

## 3 Discussion

In this work, we developed COBRA, which extends our previously published OntoVAE model with an adversarial approach that drives a separation of different covariate effects in the latent space. Thus, COBRA maintains an interpretable decoder, with neurons corresponding to TF or pathway activities, while allowing to distinguish between different views, and considering TF or pathway activities that are only related to one covariate but not the other. This is achieved by removing covariate information from the latent space with auxiliary classifiers. Simultaneously, covariate information is learned as separate embeddings. In the context of COBRA, one of the covariates is usually the celltype, which is also often the strongest confounder in a dataset, while others can represent timepoints, treatments or other cellular conditions.

We demonstrated the functionality of COBRA on two different datasets for different scenarios. First, we applied the tool on a mouse scRNA-seq dataset of ESC and MEF cells that were interferon-treated for 0 hours, 1 hour, or 6 hours to decouple the celltype effect from the treatment effect. We identified NANOG as IFN responder, and this could not be achieved by a related method. We furthermore isolated pathways that were important for either classifying the samples according to celltype or IFN stimulation. We then applied COBRA on a scRNA-seq dataset of the developing adrenal medulla to identify TFs with a general importance for differentiation independent of celltype. Finally, on the same dataset, we also demonstrated how COBRA can predict the TF activities for previously unseen celltypes.

As with any method that relies on a biological prior, biological findings made with COBRA depend on the type and the quality of this prior. They are furthermore limited to the knowledge domain covered by this prior. A further limitation of COBRA is the additivity assumptions between covariates in the latent space, neglecting possible interaction terms. However, we believe that this makes the model easier to interpret, and thus a valuable add-on to the scientific community.

## 4 Methods

*Model architecture*. COBRA is an extension of our previously published OntoVAE model [12]. OntoVAE is a variational autoencoder that replaces the decoder with a biological ontology, and is thus intrinsically interpretable as the neuron activations correspond to pathway activities. COBRA maintains this interpretability in the decoder, and allows to incorporate multi-layer priors as well as single-layer priors such as TF regulons. Additionally, COBRA implements an adversarial approach as published in [13]. This approach leads to a disentanglement of the contributions of different covariates in the latent space. Thus, the encoder learns a so-called basal state which does not contain covariate information: 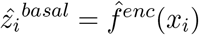. This is achieved by attaching auxiliary classifiers 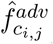 to the latent space which encourage the removal of covariate information. The auxiliary classifiers are optimized using cross-entropy: 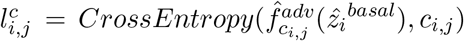, with *K* being the number of categories a covariate *c* can take (*j* = 1, …, *K*). Simultaneously, covariate embeddings 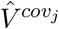 are learnt through torch.nn.Embedding layers and added to the basal state. This sum (total state) is then fed into the decoder for reconstruction: 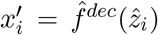. Thus, model training is an iterative process, which repeats the following two steps during training:

- The parameters of the auxiliary classifiers 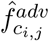 are updated to minimize 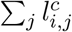
- The parameters of encoder 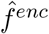, decoder 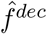, and covariate embeddings 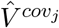 are updated to minimize 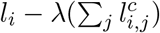

Hereby, *l*_*i*_ stands for the standard VAE loss, and *λ* is a weighting coefficient for the adversarial part of the loss.

### Model training

Throughout this manuscript, COBRA was trained using the following architecture: the latent space, a vector of length 128 which is a linear combination of basal state and different covariates, is fully connected to a hidden layer in the decoder that represents either TFs or pathways. A ReLU activation function is applied on the output of that layer, and is then sparsely connected to the genes in the final layer that are annotated to the respective TF or pathway. Training parameters such as the learning rates and the weighting coefficients of the different components of the loss function were all set to the default values of COBRA.

### Retrieval of TF and pathway activities

As in the original OntoVAE model, activities of the interpretable nodes can be retrieved with a simple function call. For COBRA, the output is a dictionary that contains the activities for each view of the trained model. The views that are independent of the dataset are the basal view where all covariate information is removed, and the total view which is a sum of the basal view and the separate covariate embeddings. The dataset specific views depend on the specified covariates and are returned as the sum of the basal view and the respective covariate embedding.

### Datasets

All datasets that were used for this manuscript are publicly available and referenced throughout the manuscript.

### Model availability

COBRA is available together with vignettes that demonstrate its usage under https://github.com/hdsu-bioquant/cobra-ai. The code for the OntoVAE model which underwent some major improvements is now also hosted in this repository.

## Acknowledgments

This work was supported by the e:Med project COMMITMENT [grant 01ZX1904D to D.D., Y.Z. and C.H.] from the German ministry for research and education (BMBF). C.R.A. and C.H. acknowledge funding through the Deutsche Forschungsgemeinschaft (DFG, German Research Foundation) – project number 272983813 (TRR 179). A.L. acknowledges financial support through the Studienstiftung des deutschen Volkes.

